# Development in the cold renders bird mitochondria more susceptible to heat stress

**DOI:** 10.1101/2024.09.12.612720

**Authors:** Maria Correia, Elisa Thoral, Elin Persson, Imen Chamkha, Eskil Elmér, Andreas Nord

## Abstract

Research on birds suggests that extreme weather events during development may have long-lasting consequences on form and function. The underlying cellular mechanisms mediating such phenotypic effects are poorly studied. We raised Japanese quail in warm (30°C) or cold (10°C) temperatures from hatching until adulthood, and then measured mitochondrial metabolism in intact blood cells at representative normothermic body temperature (41°C) and a hyperthermic temperature (45°C) that quail commonly attain when heat stressed. To investigate whether any developmental effects were reversible, half of the cold- and warm-acclimated birds were assigned to a common garden (20°C) 3 weeks before the measurements. Across groups, hyperthermia was associated with increased proton leak, but decreases in both phosphorylating respiration (where ATP is produced) and working capacity of the mitochondria. Cold-acclimated birds were more strongly affected by heat stress: the increase in proton leak was 1.6-fold higher, and the decrease in phosphorylating capacity during endogenous respiration was 1.7-fold greater, compared to warm-acclimated birds. These differences did not remain in the common-garden birds. Our study suggests that developmental cold acclimation is traded off against heat tolerance at the level of cellular metabolism, with implications for our understanding of avian responses to climate change.

## Introduction

Extreme weather events, such as heatwaves and cold spells, are predicted to increase in intensity and frequency under climate change (Roberts et al., 2021) with anticipated negative effects on the fitness and survival of wildlife (Parmesan et al., 2000; Stillman 2019). If such thermal variations are timed at sensitive windows of development (cf., Burggren 2018), evidence from mammals and birds point to potentially irreversible epigenetic modulation of morphology and physiology (e.g., Nichelmann and Tzschentke, 2002; reviewed by Nord and Giroud, 2020; Andreasson et al. 2020). For example, in fowl, increased temperature during pre- or post-natal development increases peripheral circulation (Burness et al. 2013), improves evaporative cooling capacity (Persson et al. 2024), and may reduce mortality during acute heat challenges (Arjona et al. 1988). Similarly, pre- or postnatal cold stress improves thermogenic performance, attenuates the stress response, and improves survival, during later-life cold challenges (Shinder et al., 2002, 2009). While the endocrinology of thermoregulation is reasonably well understood, also in a developmental programming perspective (reviewed by Ruuskanen et al. 2021), many aspects of the cellular responses linking temperature conditions in early life to subsequent performance during a thermal challenge remain understudied.

Thermoregulation in birds is achieved through a coordinate suite of behavioural and physiological processes that require energy in the form of adenosine triphosphate (ATP), which is produced almost exclusively via oxidative phosphorylation (OXPHOS) in the mitochondria (Lehninger et al., 1993). OXPHOS is not, however, entirely efficient as some protons leak through the inner mitochondrial membrane (LEAK respiration) via both passive and regulated processes (Jastroch et al., 2010), dissipating the proton motive force as heat at the expense of reduced ATP production. Yet, LEAK is a crucial aspect of mitochondrial function, forming the basis for non-shivering thermogenesis in mammalian brown adipose tissue (Cannon and Nedergaard, 2004) and regulating the balance between ATP production and free radical formation (Seebacher et al. 2010). It follows that mitochondrial respiration is tailored to the heat or energy requirements of individuals, increasing in the cold (e.g., Nord et al., 2021; Zheng et al., 2014) and decreasing in the warmth (Pacheco-Fuentes et al. 2023). Additionally, research on mammals show that mitochondria are amenable to epigenetic developmental programming by environmental perturbations (Gyllenhammer et al., 2020). This suggests that alterations of the mitochondrial phenotype may be a potential avenue linking variation into developmental temperature to improved thermoregulatory control at the organismal level.

In accordance with the above, studies show that elevated embryonic temperature, or the perception of heat stress through parental signalling, causes increased baseline respiration and OXPHOS in intact blood cells in juvenile birds (Udino et al. 2021, Stier et al. 2022). Moreover, a perinatal heat wave increased mitochondrial respiration in adolescent zebra finches (*Taeniopygia guttata*) through higher LEAK (Ton et al., 2021). It is unclear whether this developmental programming also impacts the thermal sensitivity of mitochondria, that is, how the mitochondria subsequently *respond* to a temperature challenge. In zebra finches, acoustic heat conditioning during embryonic development did not alter the mitochondrial response to a postnatal heat challenge (Udino et al. 2021), but another study on the same species found that constant perinatal heat exposure was associated with a reduction in LEAK upon heat acclimation in adults more than two years after initial exposure (Pacheco-Fuentes et al. 2023). To the best of our knowledge, however, no study on birds has addressed how the mitochondria themselves respond to heat stress. This data paucity is constraining the research field, not the least since most bird species respond to an acute heat challenge by (voluntary, or not) hyperthermia (McKechnie and Wolf, 2019). Therefore, studies of the thermal sensitivity of mitochondrial metabolism in a developmental temperature context could give novel insight into how animals counter extreme weather events, from cells to organisms.

To determine whether the thermal sensitivity of mitochondrial function is programmed by developmental temperature, we reared Japanese quail in simulated heatwave-(30°C) or cold snap-(10°C) conditions from hatching to reproductive maturity, after which half of the individuals from each acclimation group were transferred to a common garden at intermediate temperature (20°C). Three weeks later, we measured mitochondrial respiration in intact blood cells at both a normal body temperature (41°C) and a representative hyperthermic temperature that quail incur during heat stress (45°C) (cf. Persson et al. 2024). We expected a higher mitochondrial respiration, but lower ATP-producing efficiency, during physiological hyperthermia (Roussel & Voituron 2020; Barbe et al., 2023) due to increased LEAK, reduced OXPHOS, or both. If developmental temperature impacts thermal sensitivity, we expected these changes to be attenuated in heatwave-reared birds but exaggerated in cold-reared individuals (cf. Harada et al., 2019). Finally, if any effects of developmental temperature on thermal sensitivity reflected developmental programming, we expected phenotypic change to remain even when the thermal stimulus had been removed (i.e., in the common garden birds).

## Material and Methods

### Species and Housing

The study was performed using Japanese quail (*Coturnix japonica*), a precocial bird that is reproductively active from 6-8 weeks of age (Ottinger, 2001). Eggs were obtained from a commercial breeder (Sigvard Månsgård, Åstorp, Sweden) and were incubated at 37.5°C and 50% relative humidity (OvaEasy 190 Advance Series II, Brinsea, Weston, UK). Fifty-five of 85 eggs hatched (65%) and 49 chicks survived past 1 week post-hatching (henceforth, wph). On the hatching day (day 0), chicks were ringed with individual combinations of colour rings and were assigned to one of two housing temperature treatments (Fig. 1): Warm group (30°C; 30.02 ± 0.82°C; mean ± standard deviation) or Cold group (10°C; 9.70 ± 0.31°C). Birds were housed in groups ≤ 15 individuals, in open pens (310 × 120 × 60 cm) under a 14h:10h light:dark cycle, with food and water provided *ad libitum*. Until 3 wph, the birds were fed with Turkey Starter (25.5% protein; Lantmännen AB, Stockholm, Sweden), and from then onwards with Turkey Grower (22.5% protein; Lantmännen AB, Stockholm, Sweden). Until 2 wph, hatchlings had access to a heat lamp (35-39°C) which was programmed to allow 6 cooldown periods during daytime (10 min every 2 h until 1 wph; 30 min every 2 h until 2 wph). Birds were weighed (± 0.1 g) and wing length was measured (± 0.5 mm) weekly from 1 to 12 wph. At 9 wph, half of each thermal acclimation group were randomly allocated to a common garden at intermediate temperature (20.01 ± 0.35°C). These individuals are referred as Warm-Mild and Cold-Mild birds (Fig. 1). Final sample sizes are reported in Table S1.

**Figure 1.**
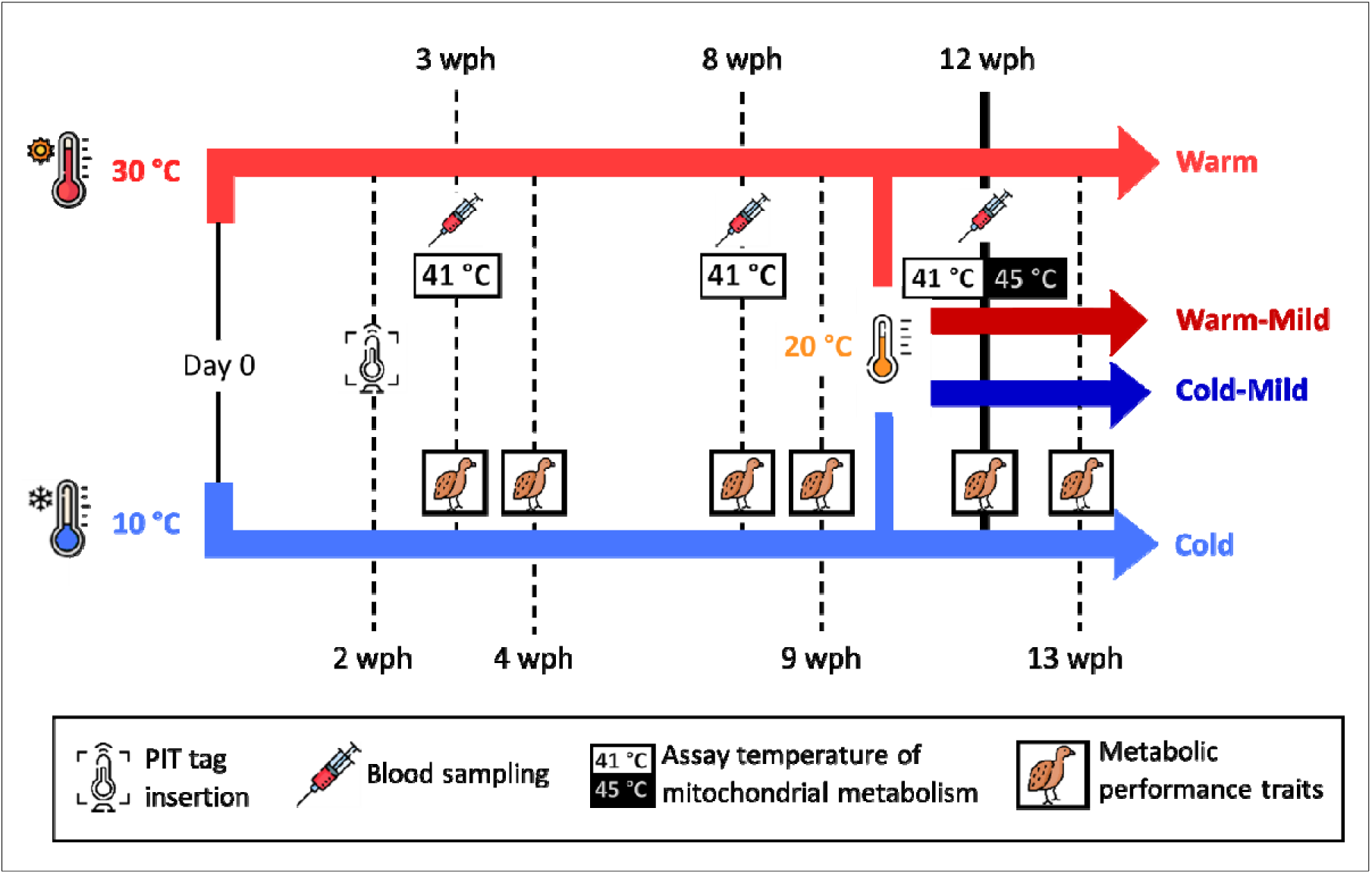
Japanese quail (*Coturnix japonica*) were reared in Warm (red, 30°C) or Cold temperature (blue, 10°C) from hatching until 9 weeks post-hatching (wph), after which half of each group was allocated to a common garden (20°C; Cold-Mild, dark blue, and Warm-Mild, dark red, groups). Birds were blood sampled and measured for metabolic performance traits at 3-4 and 8-9 wph for another study. The thermal sensitivity of mitochondrial respiration was assessed intact blood cells at 12 wph in each of a normothermic and a hypothermic assay temperature, to investigate if developmental temperature programs mitochondria function.

### Mitochondrial measurements

Blood samples (200-300 μl) were collected from the brachial vein for mitochondrial measurements at 3, 8 and 12 wph (i.e., after 3 weeks in common-garden conditions in the Warm-mild and Cold-mild groups; see Fig. 1). The birds were also measured for metabolic performance traits at 3-4, 8-9 and 12-13 wph (on the day after blood sampling) as part of a different study. To investigate if developmental temperature influenced the thermal sensitivity of mitochondria in adults, measurements of mitochondrial respiration were performed at 12 wph in 50 μl whole blood samples using an Oxygraph O2k high-resolution respirometer (Oroboros Instruments, Innsbruck, Austria) following Nord et al. (2023). The samples were measured at 41°C, which is a representative daytime body temperature of Japanese quail (Prinzinger et al., 1991; Persson et a. 2024), and at 45°C, which Japanese quail reach even during moderate heat-stress (Persson et al., 2024). We used two Oxygraphs that were calibrated at assay temperatures and air saturation oxygen level daily and changed temperature between instruments on alternate days. Mitochondrial respiration traits were assessed using intact cells in whole blood following Nord et al. (2023). Details on the protocol are presented in the ESM, and a detailed explanation of the mitochondrial traits assessed is available in Table S2.

### Statistical analyses

Statistical analyses were performed using R 4.4.1 for Windows (R Development Core Team 2024). To evaluate the impact on mitochondrial function of a simulated increase in body temperature equivalent to that experienced during heat stress, we analysed the effect of assay temperature (41°C or 45°C) on all respiration rates and flux control efficiencies (FCE’s), independently of thermal treatment, using linear mixed models (lmer in the lme4 package; Bates et al., 2015). Assay temperature, sex and body mass were used as fixed effects, and the interaction ‘assay temperature × sex’ was used as a factor. Bird identity was included as a random effect to account for repeated measurements on the same individual.

To investigate whether developmental temperature programmed the thermal sensitivity of mitochondria, we tested the mitochondrial response (i.e., the difference between traits measured at 45°C and 41°C; Δ_45°C-41°C_) in a linear model (lm in the *stats* package) with treatment, sex, treatment × sex, and body mass as explanatory variables. These models were run separately within the developmental temperature (Cold vs Warm) and common garden (Cold-Mild vs Warm-Mild) groups. Body mass was mean centred by treatment and sex in all models. Raw data collected in each assay temperature, for each developmental temperature treatment, are presented in Fig. S1.

The effects of developmental temperature (Cold or Warm) on morphological features at 12 wph (body mass, wing length) were assessed using linear models (lm in the stats package), using treatment, sex, and treatment × sex as factors.

Non-significant (*P* > 0.05) interactions were removed from the final models, but all main effects were kept. Estimates and standard errors for fixed effects were calculated using the emmeans() function in the *emmeans* package (Russell et al., 2024). Significance of random effects were analysed with ranova() function in the *lmerTest* package (Kuznetsova et al., 2017). Assumptions of normality and homogeneity of variances were determined using Shapiro-Wilk test, diagnostic plots and Levene’s test with plotresid() function in the *RVAideMemoire* package (Herve, 2023) and leveneTest() function in the *car* package (Fox and Weisberg, 2019).

## Results

### Effect of physiological hyperthermia on mitochondrial function

ROUTINE, i.e., the baseline respiration on endogenous substrates, increased significantly at the heat-stressed assay temperature (i.e., 45°C), by 1.07-fold compared to at a representative body temperature (41°C) (LMM: *p = 0.003*; Fig. 2A). This was mainly due to 1.70-fold increase in LEAK (i.e., respiration used to counteract proton leak) (*p < 0.001*), whereas OXPHOS (i.e., ATP-producing respiration) was 0.86-fold lower at 45°C (*p = 0.001*) (Fig. 2B,C). ETS (i.e., the maximum capacity of the electron transport system) was also significantly lower at a heat stressed assay temperature, by 0.79-fold (*p < 0.001*) (Fig.2D). As a result, all FCE’s (see Table S2 for definitions) indicated an overall decrease in mitochondrial efficiency during simulated heat stress (Fig. 2E-G, all *p < 0.001*). Whatever the assay temperature, females displayed significantly higher values of all mitochondrial respiration traits and FCE’s compared to males, except OXPHOS and R-L control efficiency (Table 2; Fig. S2).

**Figure 2.**
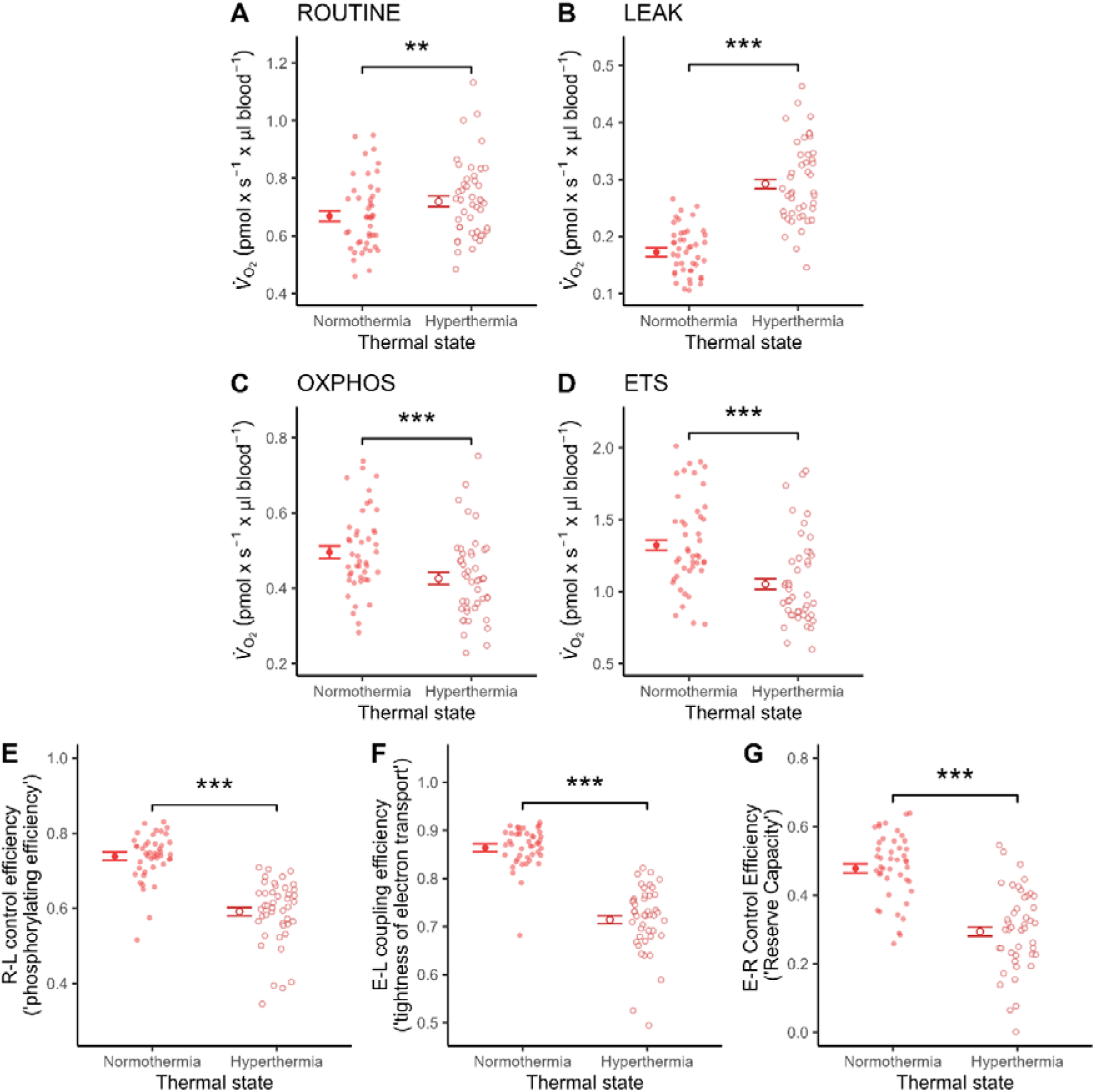
Mitochondrial respiration rates and flux control efficiencies (FCE’s) at representative normothermic (41°C) and heat-stressed (45°C) body temperatures in whole blood of 12-week-old Japanese quail. Points and error bar show estimated mean ± SE, and semi-transparent points show raw data. Asterisks (*) indicate level of significance between experimental treatments (*p* < 0.001 ‘***’; *p* < 0.01 ‘**’). *V*_O2_: oxygen consumption.

**Table 1.** Final model estimates (± standard error), degrees of freedom, test statistics and *p*-values when testing the effects of assay temperature (41°C or 45°C) on mitochondrial respiration in intact blood cells of 12-week-old Japanese quail. Significant effects (p ≤ 0.05) are printed in bold font and near-significant effects (0.05 > *p* ≤ 0.10) are printed in italics. Abbreviations: BirdID: unique identifier of each bird; df: degrees of freedom; SE: standard error; σ_ID_: standard deviation of the random effect; σ_resid_: residual standard deviation.

| Models and parameters | Estimate (± SE) | df | F | p | $\sigma_{ID}$ | $\sigma_{resid}$ |
| --- | --- | --- | --- | --- | --- | --- |
| ROUTINE (pmol O<sub>2</sub> × s<sup>-1</sup> × μl blood<sup>-1</sup>) | | | | | | |
| Assay Temperature |  | 1, 43 | 9.94 | **0.003** |  |  |
| 41°C | 0.669 (± 0.018) | | | | | |
| 45°C | 0.719 (± 0.018) | | | | | |
| Sex |  | 1, 41 | 8.59 | **0.006** |  |  |
| Female | 0.742 (± 0.022) | | | | | |
| Male | 0.645 (± 0.025) | | | | | |
| Body mass (g) |  | 1, 41 | 0.03 | 0.858 |  |  |
| BirdID |  | 1 |  | **<0.001** | 0.096 | 0.075 |
| OXPHOS (pmol O<sub>2</sub> × s<sup>-1</sup> × μl blood<sup>-1</sup>) | | | | | | |
| Assay Temperature |  | 1, 43 | 13.0 | **0.001** |  |  |
| 41°C | 0.496 (± 0.016) | | | | | |
| 45°C | 0.427 (± 0.016) | | | | | |
| Sex |  | 1, 41 | 3.54 | 0.067 |  |  |
| Body mass |  | 1, 41 | 0.35 | 0.559 |  |  |
| BirdID |  | 1 |  | **0.040** | 0.060 | 0.090 |
| LEAK (pmol O<sub>2</sub> × s<sup>-1</sup> × μl blood<sup>-1</sup>) | | | | | | |
| Assay Temperature |  | 1, 43 | 194. | **<0.001** |  |  |
| 41°C | 0.172 (± 0.008) | | | | | |
| 45°C | 0.292 (± 0.008) | | | | | |
| Sex |  | 1, 41 | 12.0 | **0.001** |  |  |
| Female | 0.256 (± 0.009) | | | | | |
| Male | 0.209 (± 0.010) | | | | | |
| Body mass |  | 1, 41 | 2.54 | 0.119 |  |  |
| BirdID |  | 1 |  | **0.004** | 0.035 | 0.040 |
| ETS (pmol O<sub>2</sub> × s<sup>-1</sup> × μl blood<sup>-1</sup>) | | | | | | |
| Assay Temperature |  | 1, 43 | 86.3 | **<0.001** |  |  |
| 41°C | 1.32 (± 0.036) | | | | | |
| 45°C | 1.05 (± 0.036) | | | | | |
| Sex |  | 1, 41 | 37.5 | **<0.001** |  |  |
| Female | 1.392 (± 0.045) | | | | | |
| Male | 0.983 (± 0.049) | | | | | |
| Body mass |  | 1, 41 | 1.04 | 0.313 |  |  |
| BirdID |  | 1 |  | **<0.001** | 0.198 | 0.137 |
| R-L Control Efficiency |  |  |  |  |  |  |
| Assay Temperature |  | 1, 43 | 94.0 | **<0.001** |  |  |

**Table 1.** (continued)
| Models and parameters | Estimate (± SE) | df | F | p | $\sigma_{ID}$ | $\sigma_{resid.}$ |
| --- | --- | --- | --- | --- | --- | --- |
| 41°C | 0.739 (± 0.011) | | | | | |
| 45°C | 0.591 (± 0.011) | | | | | |
| Sex |  | 1, 41 | 1.79 | 0.189 |  |  |
| Body mass |  | 1, 41 | 3.28 | 0.078 |  |  |
| BirdID |  | 1 |  | 0.742 | 0.016 | 0.071 |
| E-L Coupling Efficiency |  |  |  |  |  |  |
| Assay Temperature |  | 1, 43 | 199. | **<0.001** |  |  |
| 41°C | 0.864 (± 0.008) | | | | | |
| 45°C | 0.714 (± 0.008) | | | | | |
| Sex |  | 1, 41 | 4.91 | **0.032** |  |  |
| Female | 0.803 (± 0.009) | | | | | |
| Male | 0.775 (± 0.009) | | | | | |
| Body mass |  | 1, 41 | 0.99 | 0.327 |  |  |
| BirdID |  | 1 |  | 0.224 | 0.024 | 0.049 |
| E-R Control Efficiency |  |  |  |  |  |  |
| Assay Temperature |  | 1, 43 | 178. | **<0.001** |  |  |
| 41°C | 0.478 (± 0.013) | | | | | |
| 45°C | 0.294 (± 0.013) | | | | | |
| Sex |  | 1, 41 | 30.4 | **<0.001** |  |  |
| Female | 0.449 (± 0.016) | | | | | |
| Male | 0.323 (± 0.017) | | | | | |
| Body mass |  | 1, 41 | 0.72 | 0.402 |  |  |
| BirdID |  | 1 |  | **0.001** | 0.060 | 0.065 |

**Table 2.** Final model estimates (± standard error), degrees of freedom, test statistics and *p*-values when investigating the effects of developmental temperature on thermal sensitivity (Δ_45°C-41°C_) of mitochondrial respiration in intact blood cells in Japanese quail raised under Cold (10°C) or Warm (30°C) conditions. Significant effects (p ≤ 0.05) are printed in bold and near-significant effects (0.05 > p ≤ 0.10) are printed in italics. Abbreviations: df: degrees of freedom; SE: standard error.

| Models and parameters | Estimate ± SE | df | F | p |
| --- | --- | --- | --- | --- |
| ROUTINE ($\text{pmol O}_2 \times \text{s}^{-1} \times \mu\text{l blood}^{-1}$) | | | | |
| Treatment |  | 1, 17 | 0.62 | 0.443 |
| Sex |  | 1, 17 | 0.07 | 0.791 |
| Body mass |  | 1, 17 | 0.44 | 0.517 |
| OXPHOS ($\text{pmol O}_2 \times \text{s}^{-1} \times \mu\text{l blood}^{-1}$) | | | | |
| Treatment |  | 1, 16 | 3.08 | *0.097* |
| Sex |  | 1, 16 | 0.01 | 0.945 |
| Body mass |  | 1, 16 | 0.96 | 0.340 |
| LEAK ($\text{pmol O}_2 \times \text{s}^{-1} \times \mu\text{l blood}^{-1}$) | | | | |
| Treatment |  | 1, 17 | 6.27 | **0.023** |
| Cold | 0.161 (± 0.018) | | | |
| Warm | 0.098 (± 0.018) | | | |
| Sex |  | 1, 17 | 0.17 | 0.681 |
| Body mass |  | 1, 17 | 0.84 | 0.372 |
| ETS ($\text{pmol O}_2 \times \text{s}^{-1} \times \mu\text{l blood}^{-1}$) | | | | |
| Treatment |  | 1, 17 | 0.35 | 0.562 |
| Sex |  | 1, 17 | 0.56 | 0.466 |
| Body mass |  | 1, 17 | 0.01 | 0.918 |
| R-L Control Efficiency |  |  |  |  |
| Treatment |  | 1, 17 | 4.61 | **0.047** |
| Cold | -0.216 (± 0.031) | | | |
| Warm | -0.122 (± 0.030) | | | |
| Sex |  | 1, 17 | 0.05 | 0.822 |
| Body mass |  | 1, 17 | 0.86 | 0.368 |
| E-L Coupling Efficiency |  |  |  |  |
| Treatment |  | 1, 17 | 2.74 | 0.116 |
| Sex |  | 1, 17 | 0.28 | 0.603 |
| Body mass |  | 1, 17 | 0.06 | 0.813 |
| E-R Control Efficiency |  |  |  |  |
| Treatment |  | 1, 17 | 0.29 | 0.599 |
| Sex |  | 1, 17 | 0.07 | 0.799 |
| Body mass |  | 1, 17 | 3.12 | *0.096* |

### Effects of developmental temperature on the thermal sensitivity of mitochondrial function

The thermal sensitivity (i.e., the difference in respiration between 45°C and 41°C) of LEAK respiration was 1.64-fold higher in the Cold compared to the Warm birds (LM: *p = 0.023*, Fig. 3C). Likewise, the decrease of R-L control efficiency, an index of phosphorylating efficiency under endogenous cellular respiration, was significantly greater in the Cold treatment (by 1.77-fold; *p = 0.047*, Fig. 3E). There were no significant effects of developmental temperature treatment on any other respiration traits or FCE (Fig. 3). However, when removing one outlier from the dataset (Warm treatment, > 2 SD from the group mean; see Fig. S1C), the decrease of OXPHOS during simulated hyperthermia was 7.14-fold higher in the Cold (−0.15 ± 0.040 pmol O_2_ × s-1 × µl blood-1) compared to the Warm birds (−0.021 ± 0.040 pmol O_2_ × s-1 × µl blood^-1^) (*p = 0.036*, Fig. S2C). There were no significant differences in any respiration rate or FCE’s, and no covariates were significant, when comparing the Cold-Mild and Warm-Mild birds (all *p > 0.1*; Fig. S3, Table S3).

**Figure 3.**
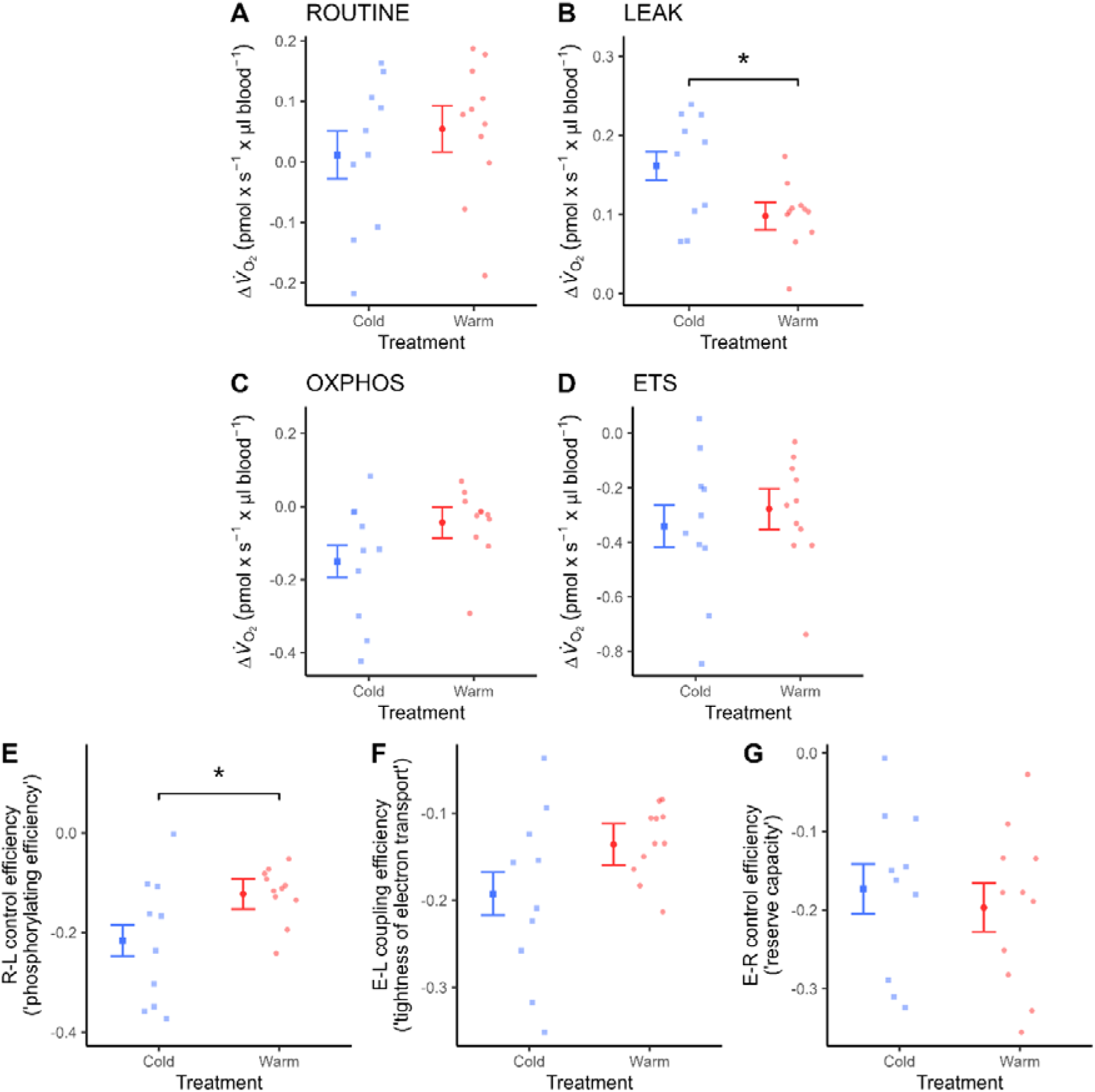
Thermal sensitivity (Δ_45°C-41°C_) of mitochondrial respiration in intact blood cells of 12-week-old Japanese quail raised in Cold (10°C) or Warm (30°C) environments from hatching onwards. Points and error bar show estimated mean ± SE, and semi-transparent points show raw data. Comparisons for which *p* < 0.05 are denoted by an asterisk (*). *V*_O2_: oxygen consumption. Δ*V*_O2_: difference in oxygen consumption recorded at 45°C and 41°C.

### Effects of developmental temperature and body mass and size

Body mass was not affected by developmental temperature (Fig. S4A, Table S4). However, Warm birds had significantly longer wings (124.13 ± 0.87 mm) compared to Cold birds (120.18 ± 0.94 mm; LM: *p = 0.006*) at 12 wph (Fig. S4B, Table S4). Moreover, females were heavier and longer-winged across treatments compared to males (Fig. S4C-D, Table S4).

## Discussion

Physiological hyperthermia simulating a body temperature that quail regularly incur during heat stress, caused significantly higher mitochondrial respiration (ROUTINE) in intact blood cells. This was driven by an increase in LEAK, which was 1.7-fold higher at 45°C compared to 41°C (Fig. 2A). ATP-producing respiration (OXPHOS), on the other hand, dropped significantly during hyperthermia (Fig. 2B). Additionally, maximum working capacity of the mitochondria (ETS) was significantly lower during heat stress. In aggregate, therefore, when the blood was exposed to hyperthermia, the phosphorylating capacity in the baseline state (R-L control efficiency), coupling efficiency in a stimulated state (E-L coupling efficiency), and mitochondrial aerobic scope (E-R control efficiency), were all significantly lower than in normothermia (Fig. 2E-G).

Acute heat stress increases the production of reactive oxygen species (ROS) and decreases antioxidant defences (reviewed by Slimen et al. 2014; Zaboli et al. 2019). Increased proton leak in a hyperthermic cell could therefore be beneficial since it reduces ROS production (Brand, 2000; Divakaruni and Brand 2011). This, in turn, may improve cell survival during a heat challenge (Omar et al. 1987). Any such benefits must, however, be leveraged against the potentially detrimental thermal effects of increased LEAK, which will add additional heat to an already heat-stressed body (Jastroch and Seebacher, 2020); a consequence that can hardly be viewed as being adaptive. Moreover, while research on mammals show that there is a regulated increase proton leak increases in heat-challenged cells (Jarmuszkiewicz et al. 2015), the expression of putative avian uncoupling proteins (avUCP, ANT) is lower or unaltered during heat stress (Mujahid et al. 2005; 2006). The precise mechanisms underpinning increased LEAK in hyperthermic cells, whether regulated by uncoupling proteins (cf. Dufour et al. 1996) or emergent from direct thermal effects acting on membrane fluidity (cf. García-Díaz et al. 2023), should be investigated in future studies.

The reductions in both OXPHOS and ETS observed in the hyperthermic condition indicate that heat stress may come at a cost of reduced ATP production, as previously demonstrated in ectotherms (Roussel & Voituron 2020). This is in contrast to a recent study of physiological hypothermia, where a corresponding reduction in assay temperature revealed no effects of temperature on neither OXPHOS nor ETS (García-Díaz et al., 2023). Thus, the thermal sensitivity of mitochondrial respiration traits appears to be nonlinear over a physiologically relevant range of body temperatures that most birds experience over the course of their annual cycle. However, the effect of hyperthermia on OXPHOS might be tissue-dependent. For example, previous research on both birds and mammals found not no significant change in phosphorylating respiration in permeabilised skeletal muscle cells during *in vitro* hyperthermia (Jarmuszkiewicz et al. 2015; Barbe et al. 2023). Alternatively, lower OXPHOS and ETS in hyperthermic blood cells could present if intracellular substrate quantity is insufficient to meet the higher metabolic requirements caused by rising temperature *per se*. It would thus be interesting to repeat the experiment using permeabilized blood cells to separate the potentially limiting effects of substrate availability (Thoral et al., 2024) from direct thermal effects acting on the mitochondrial complexes (e.g. Downs & Heckathorn 1998).

There were clear effects of developmental temperature on the thermal sensitivity of blood cell mitochondria, with cold-reared birds displaying a greater increase in LEAK and more pronounced suppression of phosphorylating efficiency (i.e., R-L control efficiency) compared to warm-acclimated birds (Fig. 3). These results could be explained if cold-induced changes in mitochondrial structure (Walter and Seebacher, 2009) or membrane fluidity (Andersson et al. 2015), rendered mitochondrial metabolism more heat-sensitive. Mechanisms aside, our work indicates that acclimation to cold temperature at the organismal level might not be compatible with maintained heat tolerance at the level of cell function. This could be relevant for understanding the apparent evolutionary trade-off between reproductive heat- and cold-tolerance that was recently demonstrated in birds (Schou et al. 2021).

Thermal sensitivity did not vary among the Cold-Mild and Warm-Mild birds, which suggests that the cold-induced mitochondrial phenotype did not reflect irreversible developmental programming. We have recently found similar reversible developmental temperature-induced plasticity at the level of organismal thermoregulation in quail (Persson et al. 2024). While reversible developmental plasticity may be more common than often appreciated (Burggren 2020), these results contrast both to studies that manipulated temperature during the embryonic period in precocial birds (Stier et al. 2022) and those subjecting altricial nestlings to an intense heatwave during the early post-hatching life (Ton et al. 2021, Pacheco-Fuentes et al. 2023), although it should be noted that neither of these experiments addressed the mitochondrial *response* to a thermal stressor. It is possible that quail, which hatch with comparatively advanced thermogenic and thermolytic capacity, are more amenable for ontogenetic programming of energy metabolism by conditions experienced before hatching.

Finally, several studies suggest that birds handle heat challenges better when exposed to warmth during early life stages (e.g. Shinder et al., 2002; Yahav and Hurwitz, 1996, Arjona et al., 1988, 1990). However, at the cellular level, growing up in a warm environment did not lead to significant improvements in thermal sensitivity of mitochondrial function compared to birds that had spent 3 weeks before the measurements in a mild temperature. This could be related to the choice of acclimation temperature, as 30°C is within the thermoneutral zone of quail (Ben-Hamo et al., 2010; Persson et al. 2024). Yet, quail reared in 30°C displayed (reversible) improved evaporative cooling capacity, a proxy for heat tolerance, compared to quail reared at mild temperature (20°C), whereas cold-acclimation (10°C) did not lead to any reductions in evaporative capacity in response to a mild heat challenge (Persson et al., 2024). A comparable mismatch between organismal and cellular responses has previously been observed in response to other environmental stressors across taxa, such as oxygen availability in goldfish *Carassius auratus* (Thoral et al., 2022). Our results, therefore, suggest that even when thermoregulatory responses are indistinguishable at the organismal level, cold-adapted birds might accrue somatic costs at a higher rate upon environmental heating, since their cells cannot keep pace. This emphasises the need to study effects of extreme weather events across levels of organisation. As a final remark, it is noteworthy that warm-acclimation had small, if any, effects on thermal sensitivity of mitochondria compared to acclimation to a mild, average temperature. This raises concerns that, much like in ectotherms (Morgan et al. 2020; Jørgensen et al. 2022), warm-blooded endotherms may have but small scope to improve the physiological tolerance of somatic function in a warming world.

## Supporting information

Electronic Supplementary Material 1

## ACKNOWLEDGEMENTS

We are grateful to Camilla Björklund and Agnieszka Czopek for assistance with animal care, and to Lars Fredriksson for technical support. Comments from Livia Saccani Hervas improved a previous version of the manuscript.

## Notes

**FUNDING:** Financial support for the project was provided by the Swedish Research Council (grant no. 2020-04686, to AN) and Stiftelsen Lunds djurskyddsfond (to AN). MC was funded by a studentship from Gyllenstiernska Krapperupsstiftelsen (grant no. KR2022-0046), EP was supported by the Crafoord foundation (grant nos. 20211007, 20221018) and ET was supported by a postdoctoral grant from the Carl Trygger Foundation (grant no. CTS21: 1173).

### Competing Interest Statement

The authors have declared no competing interest.

## References

Andersson M. N., Wang, H-L., Nord, A., Salmón, P. and Isaksson, C. (2015). Composition of physiologically important fatty acids in great tits differs between urban and rural populations on a seasonal basis. Front. Ecol. Evol. 3, 93.

Andreasson, F., Nilsson, J. Å. and Nord, A. (2020). Avian reproduction in a warming world. Front. Ecol. Evol. 8, 576331.

Arjona, A. A., Denbow, D. M. and Weaver, W. D. (1988). Effect of Heat Stress Early in Life on Mortality of Broilers Exposed to High Environmental Temperatures Just Prior to Marketing. Poult. Sci. 67, 226–231.

Arjona, A. A., Denbow, D. M. and Weaver, W. D. (1990). Neonatally-induced thermotolerance: Physiological responses. Comp. Biochem. Physiol. -- Part A Physiol. 95, 393–399.

Barbe, J., Roussel, D. and Voituron, Y. (2023). Effect of physiological hyperthermia on mitochondrial fuel selection in skeletal muscle of birds and mammals. J. Therm. Biol. 117,., 103719.

Ben-Hamo, M., Pinshow, B., Mccue, M. D., Mcwilliams, S. R. and Bauchinger, U. (2010). Comparative Biochemistry and Physiology, Part A Fasting triggers hypothermia, and ambient temperature modulates its depth in Japanese quail Coturnix japonica. Comp. Biochem. Physiol. Part A 156, 84–91.

Brand, M. D. (2000). Uncoupling to survive? The role of mitochondrial inefficiency in ageing. Exp. Gerontol. 35, 811–820.

Burggren, W. W. (2018). Developmental phenotypic plasticity helps bridge stochastic weather events associated with climate change. J. Exp. Biol. 221, jeb161984.

Burggren, W. W. (2020) Phenotypic switching resulting from developmental plasticity: Fixed or reversible? Front. Phys. 10, 1634.

Burness, G., Huard, J. R., Malcolm, E. and Tattersall, G. J. (2013). Post-hatch heat warms adult beaks: Irreversible physiological plasticity in Japanese quail. Proc. R. Soc. B Biol. Sci. 280,.

Cannon, B. and Nedergaard, J. (2004). Brown adipose tissue: Function and physiological significance. Physiol. Rev. 84, 277–359.

Divakaruni, A. S. and Brand, M. D. (2011). The regulation and physiology of mitochondrial proton leak. Physiology 3, 192–205.

Downs, C. A. and Heckathorn, S. A. (1998). The mitochondrial small heat shock protein protects NADH:ubiquinone oxidoreductase of the electron transport chain during heat stress in plants. FEBS Lett 430, 246–250.

Dufour, S., Rousse, N., Canioni, P. and Diolez, P. (1996). Top-down control analysis of temperature effect on oxidative phosphorylation. Biochem J. 314, 743–751.

Fox, J. and Weisberg, S. (2019). An R Companion to Applied Regression.

García-Díaz, C. C., Chamkha, I., Elmér, E. and Nord, A. (2023). Plasticity of mitochondrial function safeguards phosphorylating respiration during. FASEB J. 37, e22854.

Gyllenhammer, L. E., Entringer, S., Buss, C. and Wadhwa, P. D. (2020). Developmental programming of mitochondrial biology: A conceptual framework and review. Proc. R. Soc. B Biol. Sci. 287,.

Harada, A. E., Healy, T. M. and Burton, R. S. (2019). Variation in thermal tolerance and its relationship to mitochondrial function across populations of Tigriopus californicus. Front. Physiol. 10,.

Herve, M. (2023). RVAideMemoire: Testing and Plotting Procedures for Biostatistics.

Jastroch, M. and Seebacher, F. (2020). Importance of adipocyte browning in the evolution of endothermy. Philos. Trans. R. Soc. B Biol. Sci. 375,.

Jastroch, M., Divakaruni, A. S., Mookerjee, S., Treberg, J. R. and Brand, M. D. (2010). Mitochondrial proton and electron leaks. Essays Biochem 47, 53–67.

Jarmuszkiewicz, W., Woyda-Ploszczyca, A., Koziel, A., Majerczak, J. and Zoladz. J. A. (2015). Temperature controls oxidative phosphorylation and reactive oxygen species production through uncoupling in rat skeletal muscle mitochondria. *Free Rad*. Biol. Med. 83, 12–20.

Jørgensen, L. B., Ørsted, M., Malte, H., Wang, T. and Overgaard, J. (2022). Extreme escalation of heat failure rates in ectotherms with global warming. Nature 611, 93–98.

Kuznetsova, A., Brockhoff, P. B. and Christensen, R. H. B. (2017). lmerTest Package: Tests in Linear Mixed Effects Models. J. Stat. Softw. 82, 1–26.

Lehninger, A. L., Nelson, D. L. and Cox, M. M. (1993). Principles of Biochemistry. Cell Biol. Physarum Didymium 4th Ed, 393–435.

McKechnie, A. E. and Wolf, B. O. (2019). The Physiology of Heat Tolerance in Small Endotherms. Physiol. 34, 302–313.

Morgan, R., Finnøena, M. H., Jensen, H., Pélabonb, C., and Jutfelt, F. (2022). Low potential for evolutionary rescue from climate change in a tropical fish. PNAS 117, 33365–33372.

Mujahid, A., Yoshiki, Y., Akiba, Y. and Toyomizu M. (2005). Superoxide radical production in chicken skeletal muscle induced by acute heat stress. Poult. Sci. 84, 307– 314.

Mujahid, A., Yoshiki, Y., Akiba, Y. and Toyomizu M. (2005). Acute heat stress stimulates mitochondrial superoxide production in broiler skeletal muscle, possibly via downregulation of uncoupling protein content*Poult*. Sci. 85, 1259–1265.

Nichelmann, M. and Tzschentke, B. (2002). Ontogeny of thermoregulation in precocial birds. Comp. Biochem. Physiol. - A Mol. Integr. Physiol. 131, 751–763.

Nord, A. and Giroud, S. (2020). Lifelong Effects of Thermal Challenges During Development in Birds and Mammals. Front. Physiol. 11, 1–9.

Nord, A., Metcalfe, N. B., Page, J. L., Huxtable, A., McCafferty, D. J. and Dawson, N. J. (2021). Avian red blood cell mitochondria produce more heat in winter than in autumn. FASEB J. 35, e21490.

Nord, A., Chamkha, I. and Elmér, E. (2023). A whole blood approach improves speed and accuracy when measuring mitochondrial respiration in intact avian blood cells. FASEB J. 37, e22766.

Omar, R. A., Yano, S. and Kikkawa, Y. (1987). Antioxidant enzymes and survival of normal and simian virus 40-transformed mouse embryo cells after hyperthermia. Cancer Res. 47, 3473–3476.

Ottinger, M. A. (2001). Quail and other short-lived birds. Exp. Gerontol. 36, 859–868.

Pacheco-Fuentes, H., Ton., R. and Griffith, S. C. (2023). ShortL and longLterm consequences of heat exposure on mitochondrial metabolism in zebra finches (Taeniopygia castanotis). Oecologia 201, 637–648.

Parmesan, C., Root, T. L. and Willig, M. R. (2000). Impacts of extreme weather and climate on terrestrial biota. Bull. Am. Meteorol. Soc. 81, 443–450.

Persson, E., Cuív, C. Ó. and Nord, A. (2024). Thermoregulatory consequences of growing up during a heatwave or a cold snap. J. Exp. Biol 227,.

Prinzinger, R., Preßmar, A. and Schleucher, E. (1991). Body temperature in birds. Comp. Biochem. Physiol. -- Part A Physiol. 99, 499–506.

R Core Team (2024) R: A Language and Environment for Statistical Computing. R Foundation for Statistical Computing, Vienna.

Roberts, D., Pidcock, R., Chen, Y., Connors, S. and Tignor, M. (2021). IPCC Special Report.

Roussel, D. and Voituron, Y. (2020). Mitochondrial costs of being hot: effects of acute thermal change on liver bioenergetics in toads (Bufo bufo). Front. Physiol. 11,.

Ruuskanen, S., Hsu, B-Y. and Nord, A. (2021). Endocrinology of thermoregulation in birds in a changing climate. Mol. Cell. Endocrinol. 519,.

Russell, A., Lenth, V., Bolker, B., Buerkner, P., Giné-vázquez, I., Herve, M., Love, J., Singmann, H. and Lenth, M. R. V (2024). Package ‘ emmeans.’ 34, 216–221.

Schou, M. F., Bonato, M., Engelbrecht, A., Brand, Z., Svensson, E. I., Melgar, J., Muvhali, P. T., Cloete, S. W. P. and Cornwallis, C. K. (2021). Extreme temperatures compromise male and female fertility in a large desert bird. Nat Commun 12,.

Seebacher, F., Brand, M. D., Else, P. L., Guderley, H., Hulbert, A. J. and Moyes, C. D. (2010). Plasticity of oxidative metabolism in variable climates: molecular mechanisms. Physiol. Biochem. Zool. 83, 721–732.

Shinder, D., Luger, D., Rusal, M., Rzepakovsky, V., Bresler, V. and Yahav, S. (2002). Early age cold conditioning in broiler chickens (Gallus domesticus): Thermotolerance and growth responses. J. Therm. Biol. 27, 517–523.

Shinder, D., Rusal, M., Giloh, M. and Yahav, S. (2009). Effect of repetitive acute cold exposures during the last phase of broiler embryogenesis on cold resistance through the life span. Poult. Sci. 88, 636–646.

Slimen, I. B., Najar, T., Ghram, A., Dabbebi, H., Ben Mrad, M., Abdrabbah, M. (2014). Reactive oxygen species, heat stress and oxidative-induced mitochondrial damage. A review. Int. J. Hypertherm. 30, 513–523.

Stier, A., Monaghan, P. and Metcalfe, N. B. (2022). Experimental demonstration of prenatal programming of mitochondrial aerobic metabolism lasting until adulthood. Proc. R. Soc. B Biol. Sci. 289,. 20212679.

Stillman, J. H. (2019). Heat waves, the new normal: Summertime temperature extremes willimpact animals, ecosystems, andhuman communities. Physiology 34, 86–100.

Thoral, E., Farhat, E., Roussel, D., Cheng, H., Guillard, L., Pamenter, M. E., Weber, J.-M. and Teulier, L. (2022). Different patterns of chronic hypoxia lead to hierarchical adaptative mechanisms in goldfish metabolism. J. Exp. Biol. 225, 1–10.

Thoral, E., Garc, C. C. and Nord, A. (2024). The relationship between mitochondrial respiration, resting metabolic rate and blood cell count in great tits. Biol. Open 13, 1–7.

Ton, R., Stier, A., Cooper, C. E. and Griffith, S. C. (2021). Effects of heat waves during post-natal development on mitochondrial and whole body physiologyL: An experimental study in zebra finches. Front. Physiol. 12,. 661670.

Udino, E., George, J. M., McKenzie, M., Pessato, A., Crino, O. L., Buchanan, K. L. and Mariette, M. M. (2021). Prenatal acoustic programming of mitochondrial function for high temperatures in an arid-adapted bird. Proc. R. Soc. B Biol. Sci. 288,. 20211893.

Walter, I. and Seebacher, F. (2009). Endothermy in birds: Underlying molecular mechanisms. J. Exp. Biol. 212, 2328–2336.

Yahav, S. and Hurwitz, S. (1996). Induction of thermotolerance in male broiler chickens by temperature conditioning at an early age. Poult. Sci. 75, 402–406.

Zaboli, G., Huang, X., Feng, X. and Ahn, D. U. (2019). How can heat stress affect chicken meat quality? A review. Poultry Sci. 98, 1551–1556.

Zheng, W. H., Li, M., Liu, J. S., Shao, S. L. and Xu, X. J. (2014). Seasonal variation of metabolic thermogenesis in Eurasian tree sparrows (Passer montanus) over a latitudinal gradient. Physiol. Biochem. Zool. 87, 704–718.

