## Supplementary material for "Development in the cold renders bird mitochondria more susceptible to heat stress": Electronic Supplementary Material 1

**This file contains:**

- Table S1 – Sample sizes.
- Details on mitochondrial measurement techniques.
- Table S2 – Derivation and definitions of mitochondrial respiration traits.
- Figure S1 – Effects of assay temperature on mitochondrial respiration across all treatment groups.
- Figure S2 – Effects of sex on mitochondrial respiration rates and flux control efficiencies.
- Figure S3 – Thermal sensitivity of mitochondrial function in the Cold-Mild and Warm-Mild treatment groups.
- Table S3 – Model estimates and test statistics for models investigating the thermal sensitivity of mitochondrial function in the Cold-Mild and Warm-Mild groups.
- Figure S4 – Effects of thermal manipulation on final body mass and size.
- Table S4 – Model estimates and test statistics for the effects of thermal manipulation on morphometry in cold-reared and warm-reared birds.

**Table S1.** Number of individual Japanese quail (*Coturnix japonica*) used in the investigation of mitochondrial thermal sensitivity at 12 weeks post-hatching (wph). Warm-Mild and Cold-Mild refers to individuals that were transferred to a common garden at 9 wph.

| **Treatment** | **Sample size** | | |
|---|---|---|---|
| | Male | Female | Total |
| Warm (30°C) | 7 | 4 | 11 |
| Cold (10°C) | 5 | 4 | 9 |
| Warm-Mild (20°C) | 6 | 5 | 11 |
| Cold-Mild (20°C) | 6 | 6 | 12 |

### Mitochondrial measurements

Blood samples (200-300 µL) were collected by venipuncture of the brachial vein using a 1 mL heparinized syringe with a 29G needle. Samples were stored cold (10-12°C) in 2 mL $K_2$-EDTA (ethylenediaminetetraacetic acid) tubes (BD Vacutainer, Becton Dickinson AB, Plymouth, UK) until analysed 0.5-2 h later. Mitochondrial respiration of whole blood was measured in MiR05 ([0.5 mM EGTA, 3 mM $MgCl_2$, 60 mM K-lactobionate, 20 mM taurine, 10 mM $KH_2PO_4$, 20 mM HEPES, 110 mM sucrose, 1 g $l^{-1}$ free fatty acid bovine serum albumin, pH=7.1]; Gnaiger, 2020). Briefly, 50µL of whole blood were added to 1.950µL of MiR05 in the respirometer chamber calibrated at 41°C or 45°C. After closing the chamber, the baseline rate of $O_2$ consumption during respiration on endogenous substrates (i.e., "ROUTINE") was obtained for 10 min. Then, ATP synthase has been inhibited using oligomycin (1 µg $mL^{-1}$), to measure proton-leak-linked respiration (i.e., "LEAK"). Phosphorylating respiration, where ATP is produced (i.e., "OXPHOS") was defined as the difference between ROUTINE and LEAK (Gnaiger, 2020). This was followed by the determination of the maximal rate of respiration of the electron transport system (i.e., "ETS"), achieved by titration of the protonophore uncoupler Carbonyl cyanide-p-trifluoromethoxy phenylhydrazone (FCCP) until maximum (final concentration: 0.5 - 1 µM). Lastly, complex I and complex III were inhibited to determine respiration of non-mitochondrial origin (i.e., ROX respiration), using rotenone (2 µM) and antimycin (1 µg $mL^{-1}$), respectively. Since there was no further reduction in respiration when antimycin A was added on top of rotenone, ROX was defined as whichever rate was the lowest and this value was subtracted from all other respiration rates before analyses.

Three flux control efficiencies (FCE's) were calculated to provide information on mitochondrial function that is independent of any between-individual differences in mitochondrial content (Gnaiger, 2020): R-L control efficiency (endogenous respiration coupled with ATP production), E-L coupling efficiency (degree of coupling of the ETS) and E-R control efficiency (reserve capacity of the ETS).

**Table S2.** Mitochondrial respiration rates and flux control efficiencies.

| Parameter | Calculation | Definition |
|---|---|---|
| ROUTINE | Endogenous – ROX | Baseline rate of $O_2$ consumption during respiration on endogenous substrates, linked to ATP production and proton leak. |
| OXPHOS | Endogenous – Oligomycin | $O_2$ consumption used to drive ATP production. |
| LEAK | Oligomycin – ROX | $O_2$ consumption used to compensate for the proton leak. |
| ETS | FCCP – ROX | Maximum capacity of the electron transport system (ETS). |
| R-L Control Efficiency | 1 – (LEAK / ROUTINE) | Proportion of endogenous respiration used for ATP production via oxidative phosphorylation. |
| E-L Coupling Efficiency | 1 – (LEAK / ETS) | Tightness of the ETS during ATP production in a stimulated cellular state. |
| E-R Control Efficiency | 1 – (ROUTINE / ETS) | Proportion of ETS maximum working capacity used under endogenous cellular conditions. |

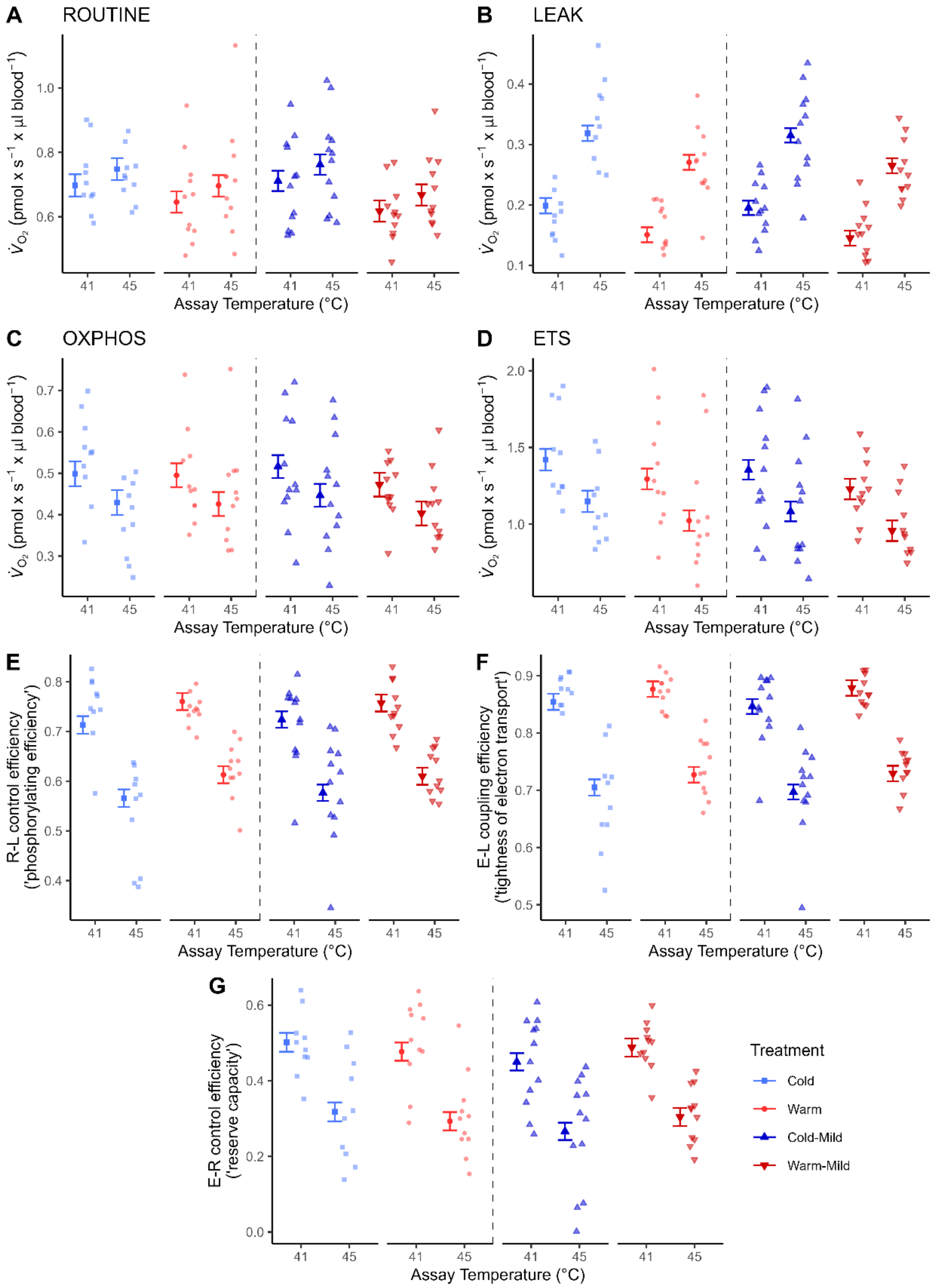


**Figure S1.** Response of mitochondrial respiration in whole blood of Japanese quail when exposed to a representative normothermic (41°C) and a representative hyperthermic (45°C) assay temperature. The birds were raised in cold (10°C) or warm (30°C) environments from hatching until 12 weeks post hatch (wph), or were transferred to a common garden (20°C; Cold-Mild, Warm-Mild) at 9 wph. Points and error bar show estimated mean ± SE, and semi-transparent points show raw data. $\dot{V}_{O_2}$: oxygen consumption.

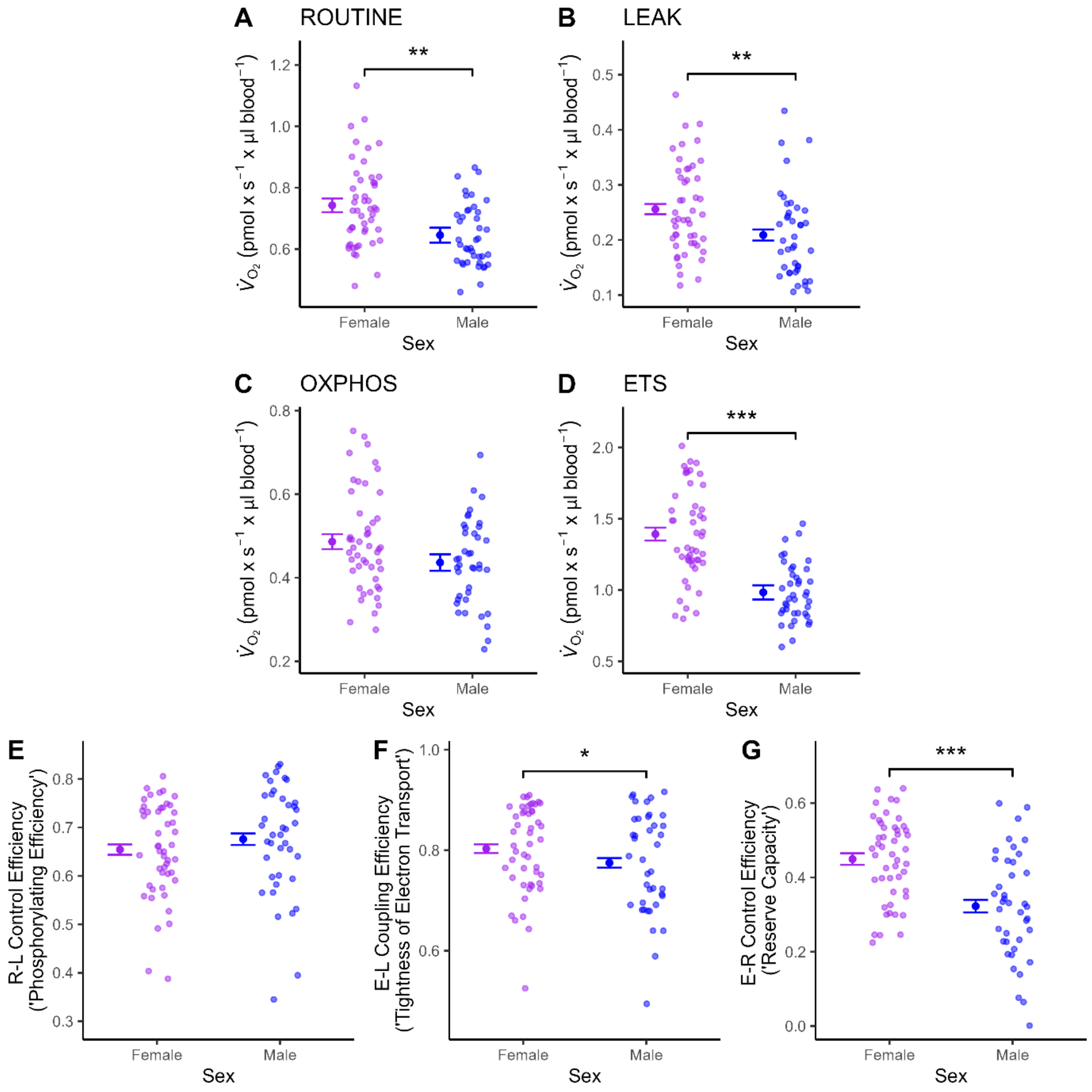


**Figure S2.** *Differences in mi*tochondrial respiration rates and flux control efficiencies (FCE's) in whole blood of 12-week-old Japanese quail between sexes across all treatments (Cold, Warm, Cold-Mild and sWarm-Mild). Asterisks (*) indicate a significant difference between the sexes ($p \leq 0.05$ '*'; $p < 0.01$ '**'; $p < 0.001$ '***'). Points and error bars show estimated mean ± SE, and semi-transparent points show raw data. $\dot{V}_{O_2}$: oxygen consumption.

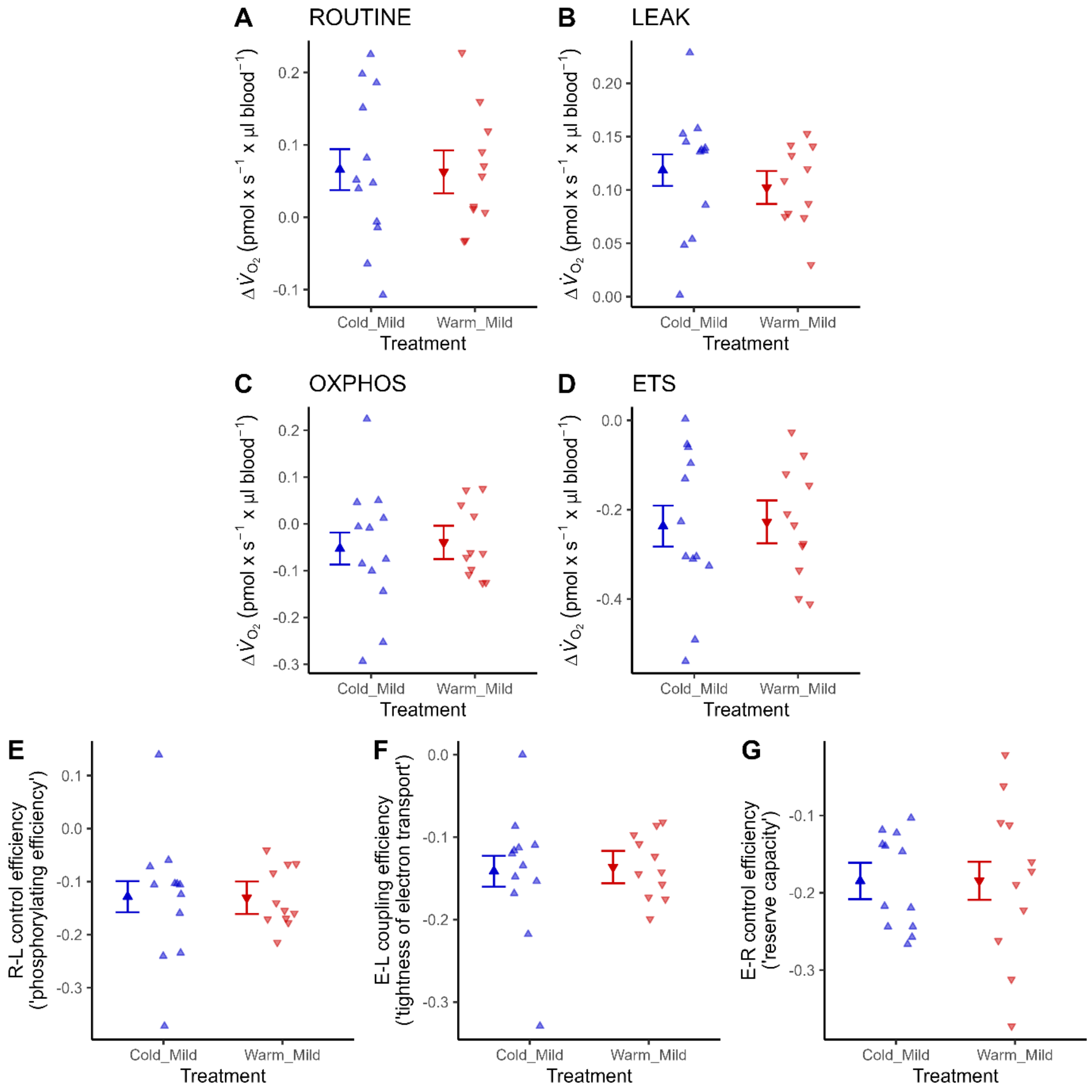


**Figure S3.** Thermal sensitivity ($\Delta_{45°C\text{-}41°C}$) of mitochondrial respiration in intact blood cells of 12-week-old Japanese quail transferred to a common garden (20°C; Cold-Mild, Warm-Mild) at 9 wph. Points and error bar show estimated mean ± SE, and semi-transparent points show raw data. $\dot{V}_{O_2}$: oxygen consumption. $\Delta\dot{V}_{O_2}$: difference in oxygen consumption recorded at 45°C and 41°C.

**Table S3.** Test statistics, degrees of freedom, and *p*-values when testing whether developmental temperature programs the thermal sensitivity of mitochondrial respiration in intact blood cells in Japanese quail. The birds were first raised under Cold (10°C) or Warm (30°C) conditions from hatching until 9-week-old, after which they were housed in a common garden at intermediate temperature (20°C) for 3 weeks prior to measurement, creating the Cold-Mild and Warm-Mild treatment groups. Abbreviations: df: degrees of freedom; SE: standard error.

| **Models and parameters** | **df** | ***F*** | ***p*** |
|---|---|---|---|
| ROUTINE (pmol $O_2$ × $s^{-1}$ × µl $blood^{-1}$) | | | |
| Treatment (Cold-Mild or Warm-Mild | 1, 19 | 0.01 | 0.941 |
| Sex | 1, 19 | 0.01 | 0.929 |
| Body mass | 1, 19 | 0.87 | 0.362 |
| OXPHOS (pmol $O_2$ × $s^{-1}$ × µl $blood^{-1}$) | | | |
| Treatment | 1, 19 | 0.07 | 0.792 |
| Sex | 1, 19 | 0.29 | 0.594 |
| Body mass | 1, 19 | 0.45 | 0.513 |
| LEAK (pmol $O_2$ × $s^{-1}$ × µl $blood^{-1}$) | | | |
| Treatment | 1, 19 | 0.58 | 0.457 |
| Sex | 1, 19 | 2.03 | 0.171 |
| Body mass | 1, 19 | 0.06 | 0.804 |
| ETS (pmol $O_2$ × $s^{-1}$ × µl $blood^{-1}$) | | | |
| Treatment | 1, 19 | 0.02 | 0.889 |
| Sex | 1, 19 | 0.21 | 0.652 |
| Body mass | 1, 19 | 0.31 | 0.587 |
| R-L Control Efficiency | | | |
| Treatment | 1, 19 | 0.00 | 0.965 |
| Sex | 1, 19 | 0.24 | 0.632 |
| Body mass | 1, 19 | 0.02 | 0.896 |
| E-L Coupling Efficiency | | | |
| Treatment | 1, 19 | 0.04 | 0.853 |
| Sex | 1, 19 | 0.48 | 0.496 |
| Body mass | 1, 19 | 0.23 | 0.638 |
| E-R Control Efficiency | | | |
| Treatment | 1, 19 | 0.00 | 0.994 |
| Sex | 1, 19 | 3.73 | 0.069 |
| Body mass | 1, 19 | 0.47 | 0.501 |

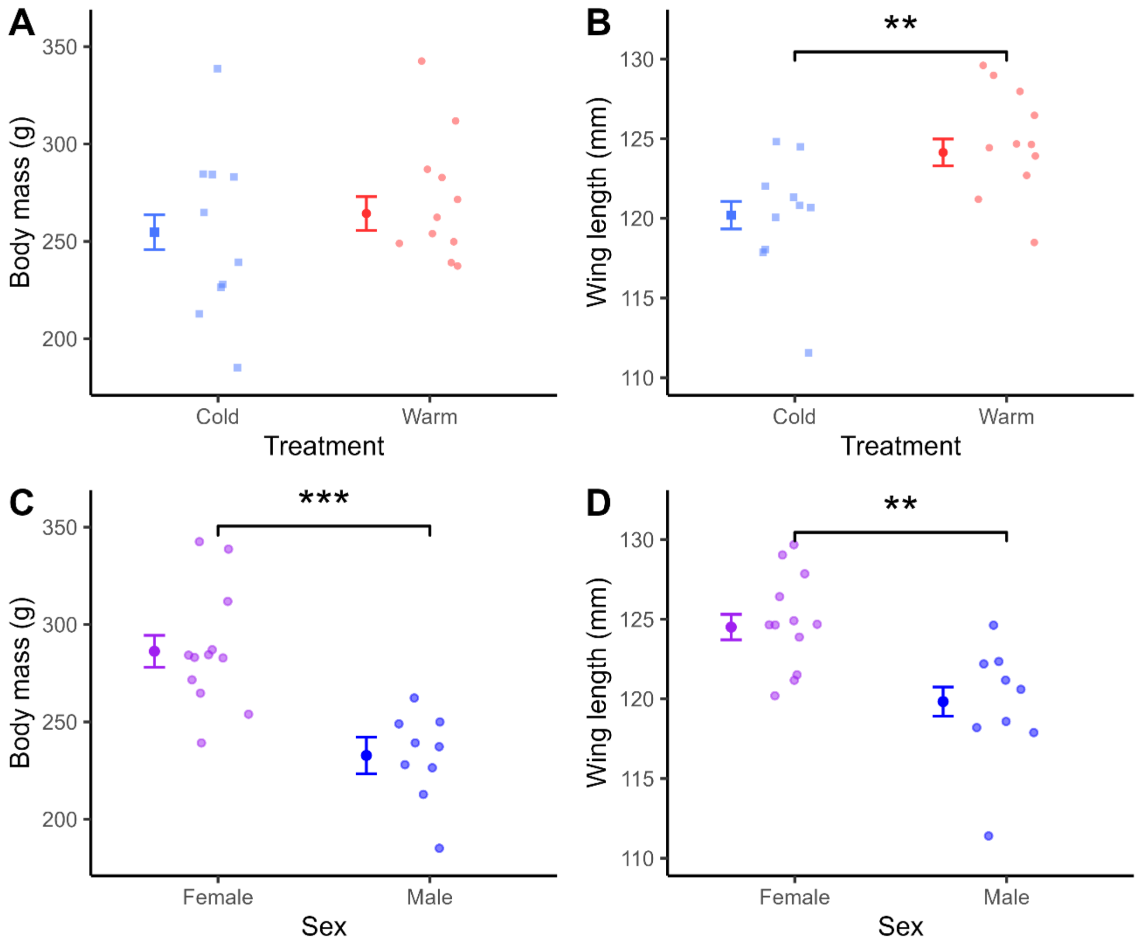


**Figure S4.** Body mass (g) and wing length (mm) of 12-week-old Japanese quail among developmental thermal environments (Cold, Warm; A-B) and across sexes, and among sexes across developmental thermal environments (C-D). Points and error bars show estimated means ± SE, and semi-transparent points show raw data. Asterisks (*) indicate level of significance between experimental treatments ($p < 0.01$ '**'; $p < 0.001$ '***').

**Table S4.** Final model estimates (± standard error), degrees of freedom, test statistics and *p*-values for tests of the effects of developmental thermal environments (Cold and Warm) on body mass and wing length. Significant effects ($p \leq 0.05$) are printed in bold. The table. Abbreviations: df: degrees of freedom; SE: standard error.

| **Models and parameters** | **Estimate ± SE** | **df** | ***F*** | ***p*** |
|---|---|---|---|---|
| Body mass | | | | |
| Treatment | | 1, 17 | 0.19 | 0.668 |
| Sex | | 1, 17 | 15.72 | **0.001** |
| Female | 286.60 (± 7.77) | | | |
| Male | 238.13 (± 9.44) | | | |
| Wing length | | | | |
| Treatment | | 1, 17 | 9.62 | **0.006** |
| Cold | 120.18 (± 0.94) | | | |
| Warm | 124.13 (± 0.87) | | | |
| Sex | | 1, 17 | 13.17 | **0.002** |
| Female | 124.50 (± 0.82) | | | |
| Male | 119.81 (± 0.99) | | | |
